# YY1-Mediated Polycomb Group Function Safeguards Hematopoietic Stem Cells from Premature Aging

**DOI:** 10.64898/2026.01.28.702390

**Authors:** Sahitya Saka, Junki P. Lee, Yinghua Wang, Peng Liu, Yunxia Liu, Courtney Hong, Ashley Kuehnl, Lixin Rui, Michael L. Atchison, Xuan Pan

## Abstract

Hematopoietic stem cells (HSCs) undergo functional decline with age, characterized by myeloid-biased differentiation, loss of quiescence, and altered metabolic homeostasis. The molecular mechanisms driving these changes remain incompletely understood. Yin Yang 1 (YY1) is a multifunctional transcription factor and mammalian Polycomb group (PcG) protein that recruits PcG complexes to specific genomic loci via its 26–amino acid REPO (Recruitment of Polycomb) domain. To define the role of YY1 PcG function in adult HSCs, we generated a conditional YY1 REPO domain knockout mouse model (*Yy1^−/ΔREPO^*). Deletion of the REPO domain led to premature HSC aging, with expansion of immunophenotypic HSCs but loss of long-term self-renewal capacity. *Yy1^−/ΔREPO^* HSCs exhibited myeloid-biased output, expansion of myeloid-primed multipotent progenitors, increased myeloid colony formation, and an elevated myeloid-to-lymphoid ratio in peripheral blood. These cells displayed reduced quiescence, elevated reactive oxygen species, increased mitochondrial oxidative capacity, and enhanced β-galactosidase activity—hallmarks of cellular aging. RNA-seq demonstrated dysregulation of gene networks governing HSC metabolism. Together, these findings establish YY1 PcG activity as a key epigenetic mechanism that preserves metabolic quiescence, sustains long-term self-renewal, and delays HSC aging. Our studies reveal a fundamental PcG-dependent epigenetic mechanism that dictate cell fate decisions and function decline during HSC aging.

## Introduction

Hematopoietic stem cells (HSCs) are undifferentiated, self-renewing, and pluripotent cells capable of generating all mature blood lineages. Maintaining a precise balance between HSC self-renewal and differentiation is critical, as disruptions lead to severe diseases such as leukaemia (1–4), lymphoma (5–7) and autoimmune disorders (8–10). However, this balance becomes increasingly dysregulated with age. While the HSC pool expands in both mice and humans during aging, this expansion fails to compensate for the functional decline that occurs. Aged HSCs exhibit a myeloid bias and reduced long-term self-renewal capacity, contributing to immune dysfunction, heightened infection susceptibility, and an increased incidence of hematologic disorders such as myelodysplastic syndrome and myelogenous leukaemia (11–13). Despite these observations, the molecular mechanisms regulating HSC differentiation and maintaining a balanced myeloid-to-lymphoid ratio during aging remain incompletely understood.

Yin Yang 1 (YY1) is a multifunctional zinc finger transcription factor and a Polycomb Group protein (PcG) that regulates both transcriptional activation and repression, in a context-dependent manner (14–16). As a key PcG protein, YY1 recruits PcG proteins to specific chromatin sites, initiating H3K27 trimethylation and stable transcriptional repression (17). In addition to its role in transcriptional regulation, YY1 promotes chromatin structural changes and initiates long-range gene regulatory DNA loops between proximal and distal promoters through homo-dimer interactions (18, 19). In murine HSPCs, YY1 physically interacts and co-localizes with chromatin structural regulator CTCF and the cohesin complex protein SMC3 throughout the genome (20). YY1 is essential for various biological processes such as early embryonic development, X-chromosome inactivation, DNA repair, and haematopoiesis (21–23). Recently, YY1 has also been identified as a key age-associated TF, with its activity significantly downregulated during HSC aging (24). Our previous work has established the fundamental role of YY1 in adult and fetal HSC biology (25, 26). Elevated YY1 levels promote HSC expansion in adult BM, whereas conditional knockout of YY1 in HSCs leads to BM failure in mice. YY1-deficient mice develop pancytopenia and die within three weeks of YY1 deletion. Furthermore, YY1-deficient HSCs fail to self-renew and lose their quiescent state (25).

Mammalian PcG proteins are key negative regulators of gene expression, binding as large complexes, such as Polycomb Repressive Complex1 (PRC1) and PRC2, to gene regulatory regions. In *Drosophila*, the recruitment of PRCs depends on the Pleiohomeotic Repressive Complex (PhoRC) (27), but a comparable mechanism in mammalian cells remains incompletely understood (28). Studies suggest that YY1 actively recruits PRC complexes in mammalian cells (29–32); however, the overlap between YY1 and PcG target genes varies across different cell types (33, 34), and other factors, such as GATA1, HIC1, REST, RUNX1, and long non-coding RNAs, have also been implicated in recruiting PRCs to specific DNA sites (35–39). These finding suggest that YY1’s PcG function may be lineage-specific. To further investigate YY1’s PcG function in haematopoiesis, we identified a 26-amino acid REPO domain, which is both necessary and sufficient for YY1 to recruit PcG proteins to specific DNA sites in *Drosophila* (29). While the YY1ΔREPO mutant can still bind DNA, activate transcription, repress transcription transiently, and interact with coregulators such as HDACs, it fails to mediate any YY1 PcG-dependent functions (29). Specifically, YY1-dependent stable transcriptional repression requires the REPO domain, which facilitates the recruitment of PcG proteins and promotes an increase in H3K27me3 at the recruitment site (17, 29, 40). Thus, a YY1 mutant lacking the REPO domain (YY1ΔREPO) serves as a powerful tool for distinguishing between YY1 PcG-dependent and PcG-independent functions. Our previous results demonstrated that YY1ΔREPO impairs Igκ chain rearrangement during B cell development, affects early T cell survival in mice and impair fetal HSC long-term self-renewal (41, 42). Furthermore, our data support that YY1’s PcG function is lineage-specific in adult haematopoiesis. In a retroviral bone marrow transplant (BMT) model, ectopic expression of YY1ΔREPO in YY1 deficient mice promotes myeloid over lymphoid development (42).

To further assess the YY1 PcG function in a physiologically relevant system, we generated a conditional YY1 REPO domain knockout mouse model (*Yy1^f/ΔREPO^ Mx1-Cre / Vav-Cre*) by genetic approach. Strikingly, deletion of the YY1 REPO domain, which mediates its PcG activity, resulted in a distinct phenotype compared with YY1 hematopoietic knockout mice and leads to premature HSC aging. *Yy1^−/ΔREPO^* HSCs are myeloid-biased and there is an expansion of myeloid-primed MPPs, increased formation of myeloid-specific colony-forming units, and an increased myeloid-to-lymphoid ratio in peripheral blood. *Yy1^−/ΔREPO^* HSCs exhibited reduced quiescence, elevated levels of intracellular reactive oxygen species (ROS), increased mitochondrial oxidative capacity, and higher β-galactosidase activity — all hallmarks of cellular aging. RNA-seq of *Yy1^−/ΔREPO^* HSCs revealed dysregulation of gene networks governing HSC metabolism, highlighting the essential role of YY1 PcG function in maintaining young metabolic homeostasis in HSCs. These findings reveal that the YY1 PcG function is essential for HSC aging by maintaining an young metabolic state. Our study uncovers a novel epigenetic mechanism through which PcG dependent mechanisms dictate cell fate decisions during the aging process in HSCs.

## Results

### Conditional deletion of the YY1 REPO domain leads to a distinct peripheral myeloid expansion

To assess the role of the YY1 REPO domain in adult hematopoiesis, we generated a *Yy1^+/ΔREPO^*mouse model using CRISPR-Cas9 gene editing (26) (**Figure 1A**). Homozygous *Yy1^ΔREPO/ΔREPO^* mice died at the early embryonic development, indicating its essential function (26). To achieve conditional deletion of the REPO domain in the hematopoietic system, *Yy1^+/ΔREPO^* mice were crossed with *Yy1^f/f^* mice to generate *Yy1^f/ΔREPO^* offspring, which were subsequently bred with *Yy1^f/+^ Mx1-Cre* or *Yy1^f/+^ Vav-Cre* mice, resulting in *Yy1 ^f/ΔREPO^ Mx1-Cre* or *Yy1^f/ΔREPO^ Vav-Cre* mice respectively (**Figure 1C**). Following Cre-mediated recombination, the wild-type *Yy1* allele is excised, leaving the ΔREPO allele as the sole functional copy (**Figure 1B** ) and the YY1ΔREPO was expressed at a comparable protein level to wild-type YY1 (**Figure 1D**). To distinguish the effects of YY1 haploinsufficiency from those of REPO domain deletion, *Yy1^+/−^*mice were used as an additional control. In contrast to YY1 knockout (KO) mice, which die within two weeks of YY1 deletion due to severe pancytopenia (25), *Yy1^f/ΔREPO^ Mx1-Cre*, *Mx1-Cre*, and *Yy1^f/+^ Mx1-Cre* mice survived for at least 200 days following poly(I:C) injection (**Figure 1E**). *Yy1^f/ΔREPO^ Mx1-Cre* mice exhibited a higher spleen-to-body weight ratio compared to both WT and YY1 KO mice (**Figure 1F**). Their liver-to-body weight ratio was similar to that of WT and heterozygous mice, but showed a slight increase relative to YY1 KO mice (**Figure 1G**). Bone marrow histology and counts revealed no evidence of aplasia in *Yy1^−/ΔREPO^*mice compared to wild-type and *Yy1^+/−^* controls (**Figures 1H, 1I & Supplemental Figure 1A**). Furthermore, complete blood counts showed no significant reductions in red blood cell numbers in *Yy1^f/ΔREPO^ Mx1-Cre* or *Vav-Cre* mice. While white blood cell counts in *Yy1^f/ΔREPO^ Mx1-Cre* mice were comparable to those of WT and heterozygous controls, *Yy1^f/ΔREPO^ Vav-Cre* mice showed a reduction relative to both control groups. Although a mild reduction in platelet count was observed in *Yy1^f/ΔREPO^ Mx1-Cre* or *Vav-Cre* mice relative to wild-type and heterozygous controls, platelet levels remained significantly higher than those in YY1 KO mice (**Figure 1J & Supplemental Figure 1B**). Compared to young HSCs, aged HSCs exhibit an enhanced capacity for myeloid differentiation and a reduced capacity for lymphoid differentiation, resulting in an increased myeloid-to-lymphoid ratio (**Figure 1K**) (43–45). Interestingly, PB analysis shows that *Yy1^f/ΔREPO^ Mx1-Cre* or *Vav-Cre* mice display a significantly higher myeloid-to-lymphoid ratio, resembling the phenotype of aged mice (**Figure 1K**). These findings support that the YY1REPO domain is essential for maintaining a balanced PB cell lineage composition. Collectively, YY1 PcG domain deleted mice exhibit distinct phenotype from YY1 KO with peripheral myeloid expansion.

**Figure 1:**
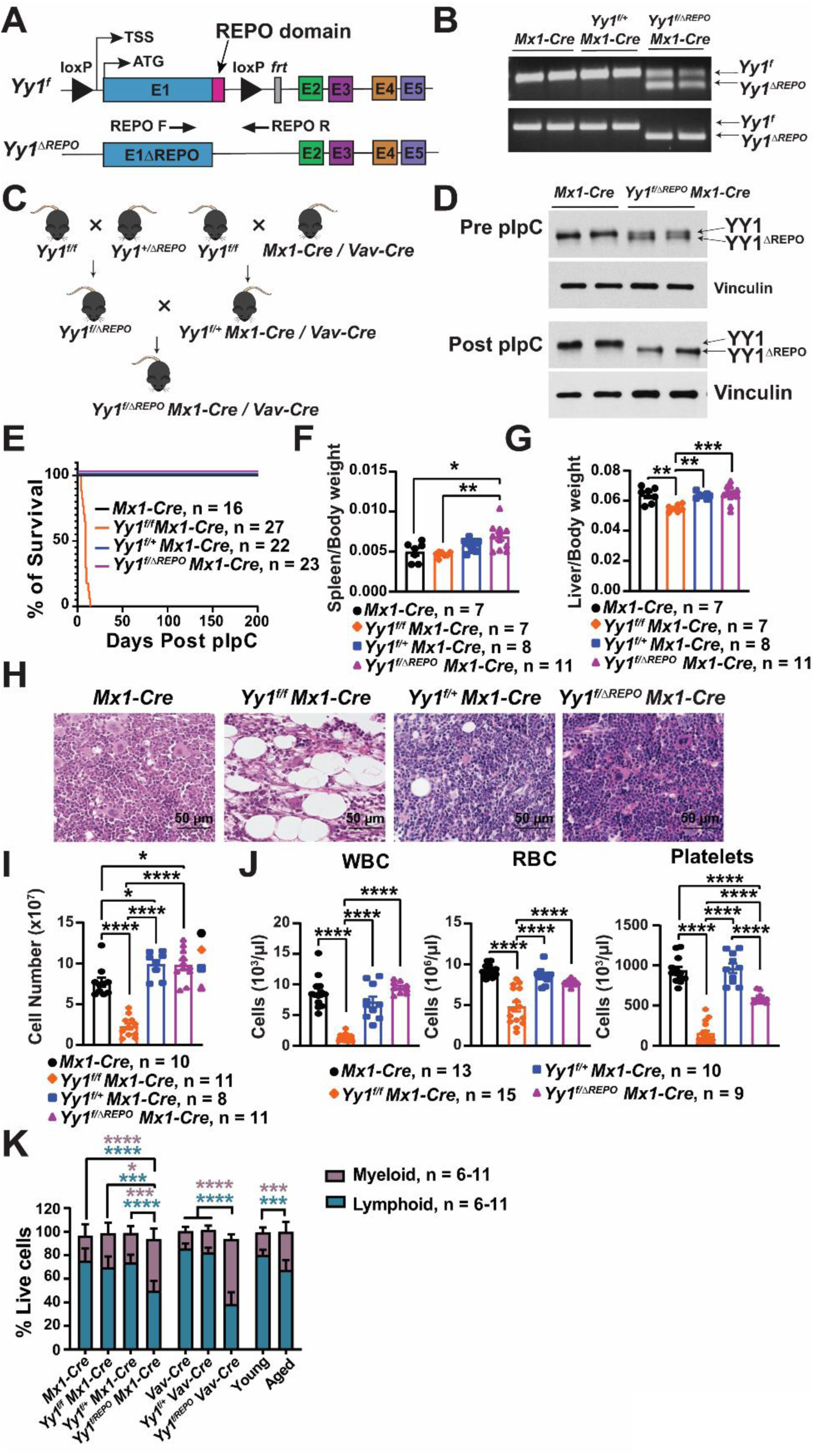
Conditional deletion of the YY1 REPO domain leads to a distinct peripheral myeloid expansion. (**A**) Schematic illustration of *Yy1^f^* and *Yy1^ΔREPO^* alleles. (**B**) PCR genotyping to show *Yy1^f^* and *Yy1^ΔREPO^* alleles. (**C**) Diagram illustration of breeding strategy to generate *Yy1^f/ΔREPO^ Mx1-Cre / Vav-Cre* mice. (**D**) Western blot showing WT YY1 (upper band) and YY1ΔREPO (lower band) before and after pI-pC injections. (**E**) Kaplan-Meier survival curve of *Mx1-Cre, Yy1^f/f^ Mx1-Cre, Yy1^f/+^ Mx1-Cre, and Yy1^f/ΔREPO^ Mx1-Cre* mice. (**F**) Quantification of spleen/body weight ratios. (**G**) Quantification of liver/body weight ratios. (**H**) BM histology. (**I**) Total BM numbers. (**J**) Complete blood count analysis. (**K**) Quantification of percentage of myeloid and lymphoid cells in PB. N represents the number of mice; Data are presented as means ± SEM; *P < 0.05, **P < 0.01, ***P < 0.001, ****P < 0.0001.

### The YY1 REPO domain deletion leads to expansion of myeloid-biased HSCs

Our previous study show that loss of YY1 led to a marked reduction in adult BM HSC percentages and numbers (25). To further investigate YY1 PcG regulation of adult HSCs, we analyzed HSC and progenitor populations in WT, *Yy1^+/−^* and *Yy1^−/ΔREPO^*mice by flow cytometry. In contrast to YY1 KO mice, which displayed a significant depletion of HSCs, *Yy1^f/ΔREPO^ Mx1-Cre* mice exhibited a pronounced increase in both percentages and absolute numbers of LSK (Lin^−^Sca1^+^c-kit^+^), LT-HSC (Lin^−^Sca1^+^c-kit^+^CD150^+^CD48^−^), ST-HSC (Lin^−^Sca1^+^c-kit^+^CD150^−^CD48^−^) and multipotent progenitor (MPP) (Lin^−^Sca1^+^c-kit^+^CD150^−^CD48^+^) populations compared with *Yy1^f/+^ Mx1-Cre* and *Mx1-Cre* controls (**Figures 2A** & **2B**). Using *Vav-Cre* as a second hematopoietic-specific deletion model, we consistently observed expansion of phenotypic LT-HSCs and ST-HSCs in *Yy1^f/ΔREPO^ Vav-cre* mice relative to *Yy1^f/+^ Vav-Cre* and *Vav-Cre* controls (**Supplemental Figure 1C**).

**Figure 2:**
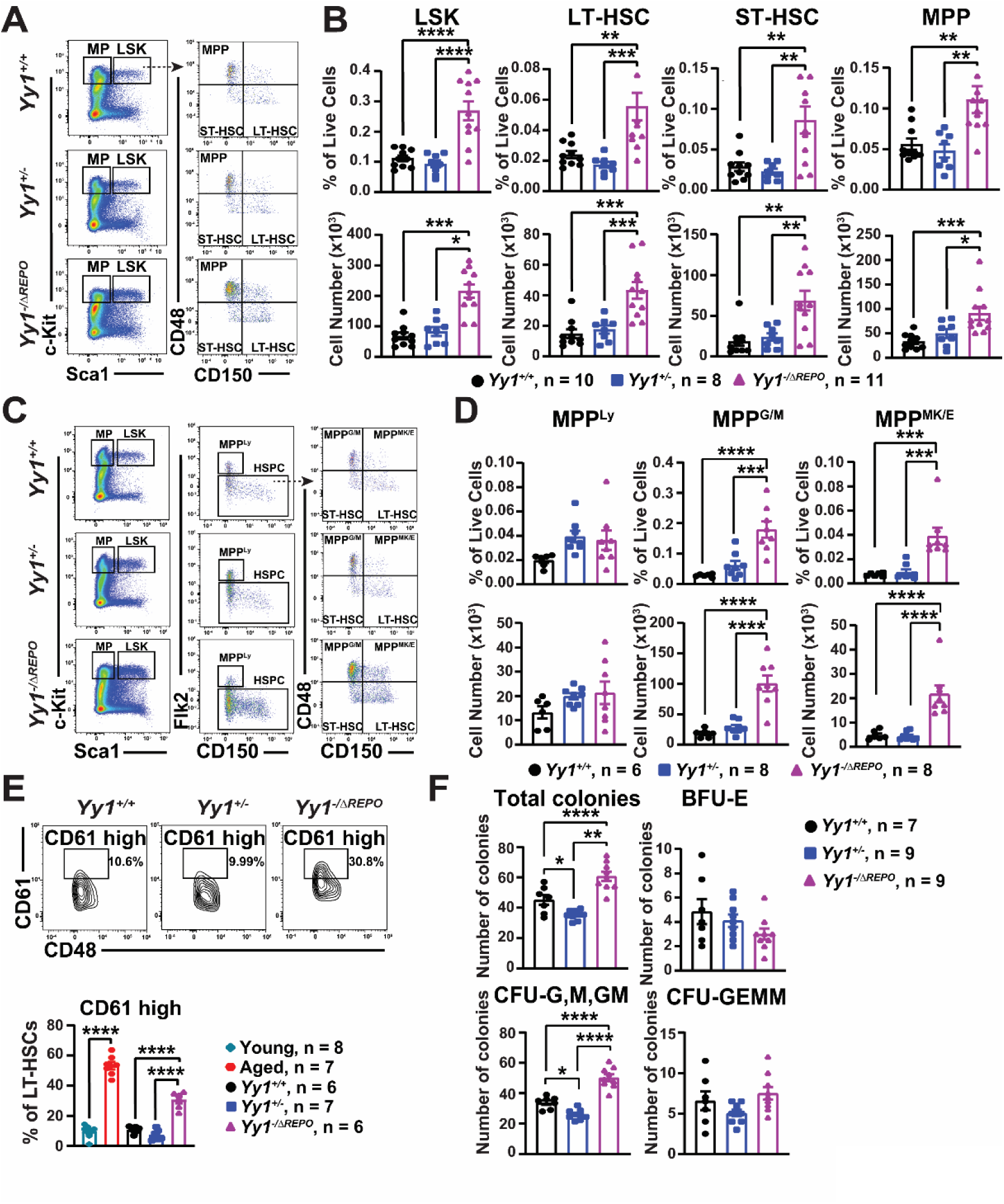
The YY1 REPO domain deletion leads to expansion of myeloid-biased HSCs. (**A**) Representative flow gating strategy for LSK (Lin^−^Sca1^+^c-kit^+^), MP (Lin^−^Sca1^−^c-kit^+^), LT-HSC (Lin^−^Sca1^+^c-kit^+^CD150^+^CD48^−^), ST-HSC (Lin^−^Sca1^+^c-kit^+^CD150^−^CD48^−^), and MPP (Lin^−^ Sca1^+^c-kit^+^CD150^−^CD48^+^) populations in BM. (**B**) Quantification of percentage and absolute number of LSK, LT-HSC, ST-HSC, and MPP populations. (**C**) Representative gating strategy for MPP^Ly^ (Lin^−^Sca1^+^c-Kit^+^Flk2^+^CD150^−^CD48^+^), MPP^G/M^ (Lin^−^Sca1^+^c-Kit^+^Flk2^−^CD150^−^CD48^+^) and MPP^MK/E^ (Lin^−^Sca1^+^c-Kit^+^Flk2^−^CD150^+^CD48^+^) populations in BM. (**D**) Quantification of percentage and absolute numbers of MPP^Ly^, MPP^G/M^ and MPP^MK/E^ populations. (**E**) Representative gating strategy and quantification of percentage of CD61 high LT-HSCs. (**F**) Colony formation assay from total BM cells. N represents the number of mice; Data are presented as means ± SEM; *P < 0.05, **P < 0.01, ***P < 0.001, ****P < 0.0001.

As *Yy1^−/ΔREPO^* mice exhibited an increased myeloid-to-lymphoid ratio in peripheral blood, we next assessed whether *Yy1^−/ΔREPO^*HSCs were lineage-biased. We quantified lineage-restricted MPP subset where MPP^Ly^ (Lin^−^Sca1^+^c-Kit^+^Flk2^+^CD150^−^CD48^+^) are lymphoid-biased, and MPP^G/M^ (Lin^−^Sca1^+^c-Kit^+^Flk2^−^CD150^−^CD48^+^) and MPP^MK/E^ (Lin^−^Sca1^+^c-Kit^+^Flk2^−^CD150^+^CD48^+^) are myeloid poised (46) (**Figure 2C**). *Yy1^−/ΔREPO^*mice showed a significant increase in both percentages and absolute numbers of MPP^G/M^ and MPP^MK/E^ populations compared with *Yy1^+/+^*and *Yy1^+/−^* controls, with no change in MPP^Ly^ frequency in any group (**Figure 2D**). Given that myeloid-biased HSC expansion during aging can be marked by increased CD61 expression (**Figure 2E**) (47), we analysed the expression of CD61 on LT-HSCs in *Yy1^−/ΔREPO^* and control mice. Notably, even in young mice, *Yy1^−/ΔREPO^* LT-HSCs displayed an elevated proportion of CD61 high cells compared with WT and *Yy1^+/−^*mice (**Figure 2E**). Functionally, total BM from *Yy1^−/ΔREPO^* mice generated more myeloid colonies including CFU-G, CFU-M, CFU-GM than WT and heterozygous controls (**Figure 2F**). Collectively, these results demonstrate that deletion of the YY1 REPO domain drives expansion of phenotypic HSC and progenitor populations, preferentially promoting myeloid-biased differentiation.

### *Yy1^−/ΔREPO^* HSCs failed to long-term self-renew

Aged HSCs have reduced long-term self-renewal capacity (11, 13, 44). We next assessed long-term self-renewal capacity of *Yy1^−/ΔREPO^* HSCs. LT-HSCs from *Mx1-Cre*, *Yy1^f/+^ Mx1-Cre* and *Yy1^f/ΔREPO^ Mx1-Cre* mice were sorted by FACS and mixed with freshly isolated WT CD45.1 BM cells and then transplanted into lethally irradiated CD45.1 recipient mice. Endogenous YY1 was deleted after engraftment was established at 4 weeks post HSC transplantation by injection of pI-pC. Peripheral blood chimerism from primary HSC transplant and consecutive secondary transplant showed significant reduction in donor derived percentages in mice transplanted with *Yy1^f/ΔREPO^ Mx1-Cre* HSCs compared to mice transplanted with *Mx1-Cre* and *Yy1^f/+^ Mx1-Cre* HSCs (**Figures 3A** & **3B**). BM HSPC analysis at 20 weeks post primary HSC transplant (**Figure 3C**) revealed significant reduction in donor derived percentages of LSK, LT-HSC, MPP in mice transplanted with *Yy1^f/ΔREPO^ Mx1-Cre* LT-HSC compared to *Mx1-Cre* control (**Figure 3D**) although no significant difference was detected between mice transplanted with *Yy1^f/ΔREPO^ Mx1-Cre* and *Yy1^f/+^Mx1-Cre* LT-HSC. In addition, at 16 weeks post-secondary transplant, mice transplanted with *Yy1^f/ΔREPO^ Mx1-Cre* cells had significant reduction of donor chimerism in HSPCs compared with both *Yy1^+/−^* and WT (**Figure 3E**). Our result supports that the YY1 REPO domain is required for HSC long-term self-renew. While YY1 heterozygosity leads to some degree of reduction of HSC self-renewal function, *Yy1^−/ΔREPO^* HSCs failed to reconstitute blood upon secondary transplantation. Our data support that the REPO domain is essential for HSC autonomous function and the YY1 REPO domain deletion leads to expansion of phenotypic LT-HSCs with defective long-term self-renewal functions.

**Figure 3:**
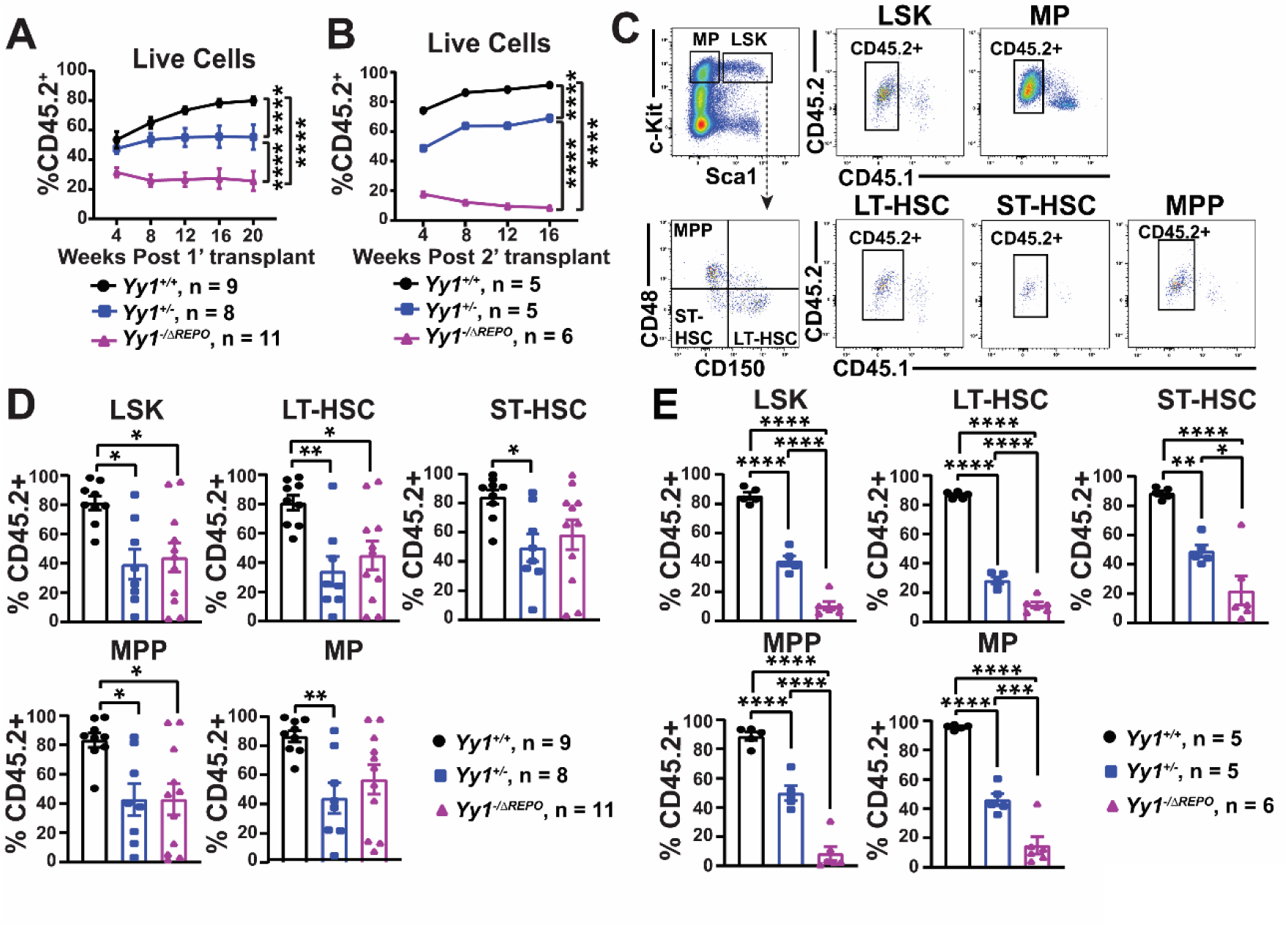
*Yy1^−/ΔREPO^* HSCs failed to long-term self-renew. (**A**) Evaluation of donor derived contribution in PB at 4-, 8-, 12-, 16- and 20-weeks post primary HSC transplantation. (**B**) Evaluation of donor derived contribution in peripheral blood at 4-, 8-, 12-, and 16-weeks post 2^nd^ transplantation. (**C**) Representative gating strategy for donor derived BM LSK, MP, LT-HSC, ST-HSC, MPP populations. (**D**) Donor derived contribution in BM LSK, LT-HSC, ST-HSC, MPP, and MP populations post primary HSC transplant. (**E**) Donor derived contribution in BM LSK, LT-HSC, ST-HSC, MPP, and MP populations post 2^nd^ transplant. N represents the number of mice; Data are presented as means ± SEM; *P < 0.05, **P < 0.01, ***P < 0.001, ****P < 0.0001.

### Genetic network governing cell metabolism were deregulated in *Yy1^−/ΔREPO^* HSCs

To further elucidate the mechanisms underlying YY1-mediated PcG regulation of adult hematopoiesis, bulk RNA-seq was performed with HSCs sorted from *Yy1^+/+^*, *Yy1^+/−^* and *Yy1^− /ΔREPO^*BM. Principal component analysis (PCA) show that *Yy1^−/ΔREPO^* samples are well separated from both *Yy1^+/+^* and *Yy1^+/−^* cohorts (**Figure 4A**). There were 231 downregulated and 126 upregulated genes detected in *Yy1^−/ΔREPO^* HSCs compared to *Yy1^+/+^* HSCs (**Figure 4B**) and 68 downregulated and 11 upregulated genes detected in *Yy1^−/ΔREPO^*HSCs compared to *Yy1^+/−^* HSCs (**Figure 4C**). Gene set enrichment analysis (GSEA) comparing *Yy1^−/ΔREPO^* to WT or *Yy1^+/−^* showed enrichment of pathways involved in metabolism, citric acid TCA cycle and respiratory electron transport, and respiratory electron transport ATP synthesis (**Figures 4 D-F**). Genes involved in cell metabolism, mitochondria function and ageing were deregulated in *Yy1^−/ΔREPO^*HSCs compared to WT and *Yy1^+/−^* (**Figure 4G**). Our data support that deletion of YY1 PcG function leads to altered HSC metabolic and aging gene signature.

**Figure 4:**
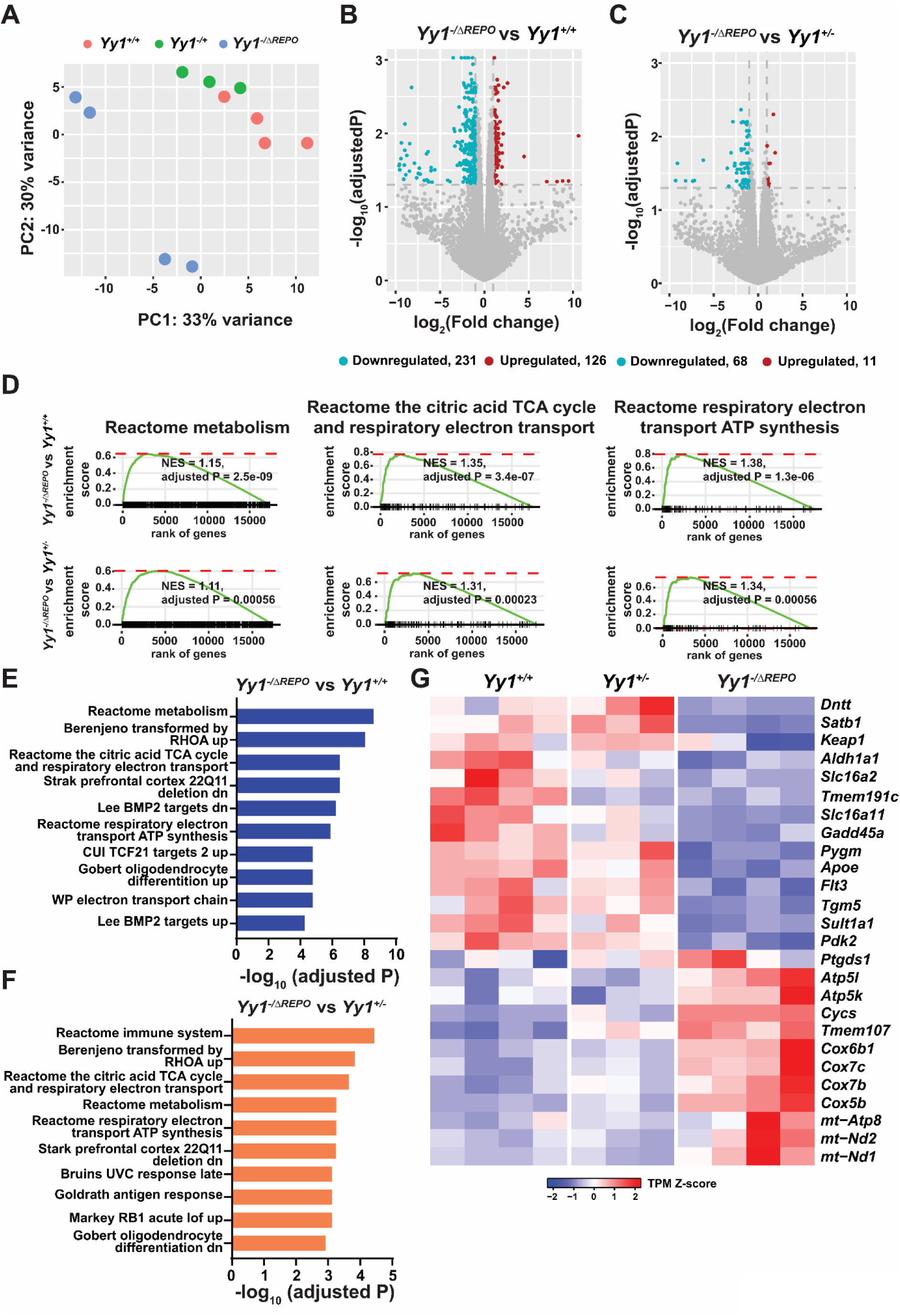
Genetic network governing cell metabolism were deregulated in *Yy1^−/ΔREPO^* HSCs. (**A**) PCA plot of RNAseq from sorted *Yy1^+/+^*, *Yy1^+/−^, and Yy1^−/ΔREPO^* HSCs. (**B**) Volcano plot showing the deregulated genes in *Yy1^−/ΔREPO^* HSCs compared to *Yy1^+/+^* HSCs. (**C**) Volcano plot showing the deregulated genes in *Yy1^−/ΔREPO^* HSCs compared to *Yy1^+/−^*HSCs. (**D**) GSEA analysis. Enriched biological processes are shown with corresponding adjusted P values and normalized enrichment score (NES). (**E**) Gene set enrichment analysis of top 10 deregulated pathways in *Yy1^−/ΔREPO^* HSCs compared to *Yy1^+/+^*. (**F**) GSEA of top 10 deregulated pathways in *Yy1^−/ΔREPO^* HSCs compared to *Yy1^+/−^*. (**G**) Heat map depicting selected upregulated and downregulated genes involved in metabolism, mitochondrial function and ageing.

### *Yy1^−/ΔREPO^* HSCs exhibit aged metabolism

It is known that aged HSCs are hyperproliferative with loss of inert metabolism (43, 48–52). We next assessed proliferation and metabolism of *Yy1^−/ΔREPO^* HSCs. Cell cycle G0 (Ki-67^−^DAPI^−^), G1 (Ki-67^+^DAPI^−^) and S/G2/M (Ki67^+^DAPI^+^) phases were defined in LT-HSC and ST-HSC populations by using Ki67/DAPI staining (**Figure 5A**). In *Yy1^f/ΔREPO^ Mx1-Cre* mice, there was a decrease in the percentage of cells in the G0 phase and an increase in the percentage of cells in the G1 phase in LT-HSC and ST-HSC compartments compared with *Mx1-Cre* and *Yy1^f/+^ Mx1-Cre* mice (**Figure 5B**). Similar results were observed in *Vav-Cre* specific deletion model (**Figure 5C**). This indicates that the YY1 REPO domain deletion led to loss of HSC quiescence and increased proliferation. We next assessed mitochondrial function and intracellular reactive oxygen species (ROS) levels, a key marker of cellular metabolic dysregulation and ageing (49) in *Yy1^+/+^*, *Yy1^+/−^* and *Yy1^−/ΔREPO^* HSPCs. Seahorse XF Mito Stress Tests on Lin⁻c-Kit⁺ cells showed that *Yy1^−/ΔREPO^* HSPCs had increased basal and maximal respiration as well as increased ATP production compared to WT and *Yy1^+/−^*HSPCs (**Figure 5D**). Similar to aged HSCs, *Yy1^−/ΔREPO^* LT-HSCs had significantly increased ROS levels compared to WT and *Yy1^+/−^* HSCs and *Yy1^−/ΔREPO^* ST-HSCs had increased ROS levels compared with WT cells (**Figure 5E**). We next assessed β-Galactosidase activity in *Yy1^−/ΔREPO^* HSCs as an indication for senescence. YY1 REPO domain deleted LT-&ST-HSCs had significantly increased β-Galactosidase activity compared to controls (**Figure 5F**) which further confirmed the ageing characteristics of REPO domain deleted HSCs.

**Figure 5:**
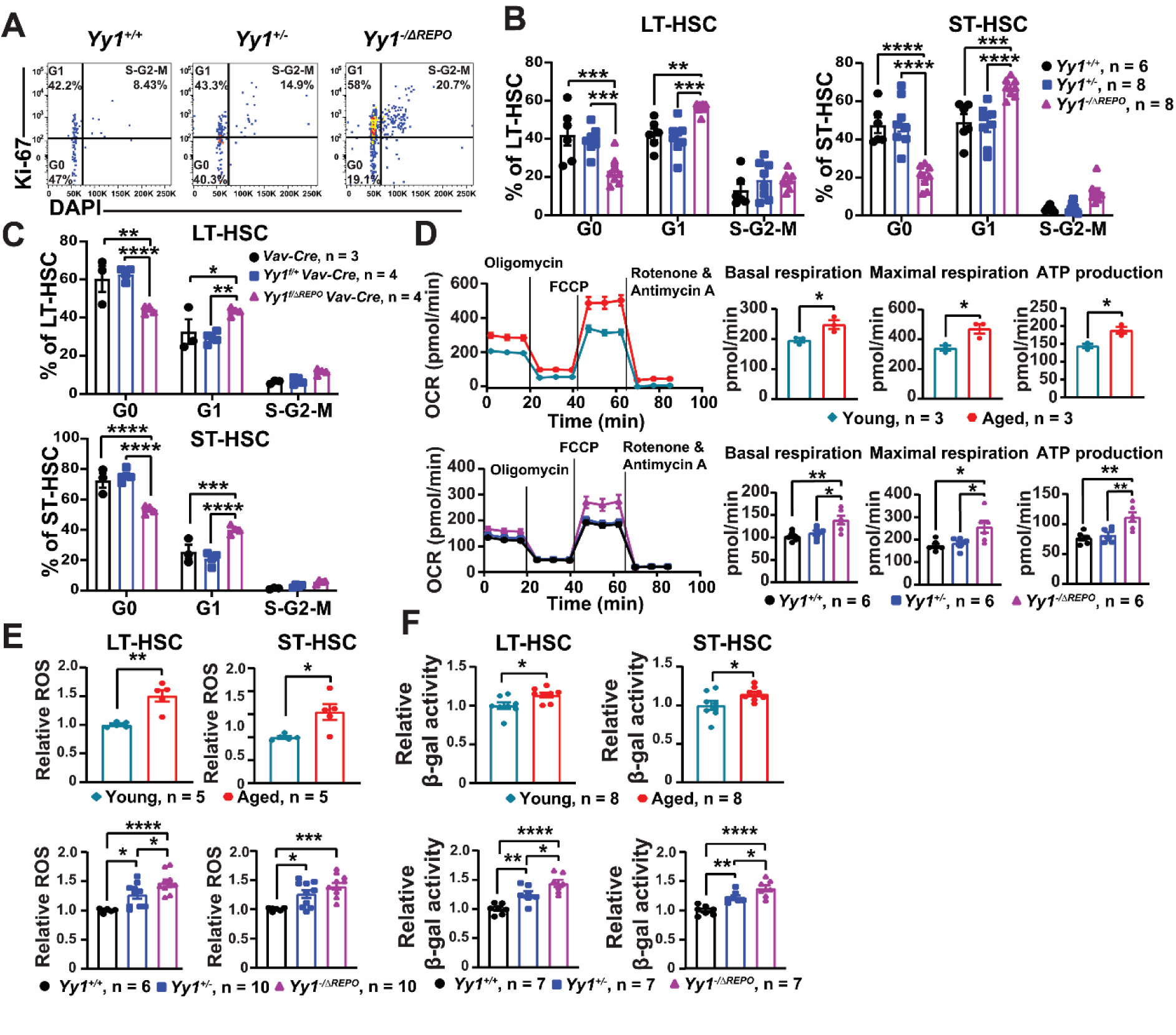
YY1 REPO domain deletion in hematopoietic system leads to aged metabolism. (**A**) Representative gating strategy for the Ki67/DAPI proliferation assay. LT-HSCs were gated for G0 (Ki67^−^DAPI^−^), G1 (Ki67^+^DAPI^−^) and S/G2/M (Ki67^+^DAPI^+^) phases. (**B**) Quantification of percentage of LT- & ST-HSCs in G0, G1, S-G2-M phases in *Mx1-Cre*, *Yy1^f/+^ Mx1-Cre* and *Yy1^f/ΔREPO^ Mx1-Cre* mice post pIpC. (**C**) Quantification of percentage of LT- & ST-HSCs in G0, G1, S-G2-M phases in *Vav-Cre*, *Yy1^f/+^ Vav-Cre* and *Yy1^f/ΔREPO^ Vav-Cre* mice. (**D**) Seahorse assay to detect OCR in LK cells. (**E**) Quantification of intracellular ROS levels in LT-& ST-HSCs. (**F**) Quantification of β-galactosidase activity in LT-& ST-HSCs. N represents the number of mice; Data are presented as means ± SEM; *P < 0.05, **P < 0.01, ***P < 0.001, ****P < 0.0001.

## Discussion

Aging of the hematopoietic system is characterized by expansion of phenotypic HSCs accompanied by functional decline, myeloid-biased differentiation, loss of quiescence, and metabolic dysregulation. While epigenetic mechanisms have long been implicated in HSC aging, the specific factors that preserve youthful HSC function remain incompletely defined. In this study, we identify the polycomb recruitment function of YY1, mediated by its REPO domain, as a critical epigenetic mechanism that restrains premature HSC aging. Using a conditional YY1 REPO domain deletion model, we demonstrate that loss of YY1 PcG activity leads to expansion of immunophenotypic HSCs with impaired long-term self-renewal, myeloid-biased differentiation, metabolic activation, and hallmarks of cellular aging.

A key finding of this study is the striking phenotypic divergence between complete YY1 loss and selective deletion of its REPO domain. Whereas YY1 knockout causes rapid bone marrow failure and lethality, *Yy1^−/ΔREPO^* mice maintain gross hematopoiesis yet develop a progressive aging-like phenotype marked by myeloid expansion and defective stem cell function. This uncoupling of YY1 PcG-dependent and PcG-independent functions highlights the modular nature of YY1 activity and underscores the specific importance of PcG recruitment in sustaining long-term HSC integrity. Our data further demonstrate that YY1 haploinsufficiency alone is insufficient to recapitulate the full phenotype, emphasizing that loss of PcG-mediated repression—not reduced YY1 dosage per se—drives premature HSC aging.

Despite a robust expansion of phenotypic LT-HSCs and ST-HSCs, *Yy1^−/ΔREPO^*HSCs exhibit severely compromised long-term self-renewal in serial transplantation assays. This paradox mirrors a defining feature of physiologically aged HSCs, in which numerical expansion masks intrinsic functional decline. The increased proportion of CD61^high^ LT-HSCs, expansion of myeloid-primed MPP subsets, enhanced myeloid colony formation, and elevated myeloid-to-lymphoid ratios in peripheral blood collectively indicate that YY1 PcG function is essential for preserving balanced lineage output. These findings place YY1 PcG activity among a growing group of epigenetic regulators, including PRC1 and PRC2 components that constrain myeloid skewing and maintain stem cell fidelity during aging.

Mechanistically, our transcriptomic and functional analyses reveal that YY1 PcG activity is a key regulator of HSC metabolic homeostasis. RNA-seq analysis identified significant dysregulation of pathways involved in mitochondrial function, oxidative phosphorylation, and cellular metabolism in *Yy1^−/ΔREPO^* HSCs. Functionally, these transcriptional changes translate into increased mitochondrial respiration, elevated ATP production, and heightened ROS levels—features that closely resemble aged HSCs. The loss of quiescence, as evidenced by reduced G0 occupancy and increased cycling, further supports the notion that YY1 PcG function actively enforces a metabolically restrained, dormant stem cell state. Importantly, increased β-galactosidase activity in YY1 REPO-deficient HSCs indicates activation of senescence-associated programs, suggesting that metabolic activation may directly contribute to irreversible functional decline.

These findings support a model in which YY1-mediated PcG recruitment represses metabolic and aging-associated gene networks to preserve a youthful HSC state. By maintaining stable transcriptional repression through H3K27me3 deposition, YY1 PcG activity likely buffers HSCs against metabolic stress and proliferative exhaustion. This function may be particularly critical in aging, where cumulative environmental and intrinsic stressors challenge stem cell homeostasis. Notably, YY1 has been identified as an age-associated transcription factor whose activity declines in aged HSCs, raising the possibility that reduced YY1 PcG function contributes causally to physiological HSC aging.

Our study also reinforces the concept that PcG regulation is highly context- and lineage-specific. Although YY1 is broadly expressed and multifunctional, selective disruption of its PcG recruitment function produces a phenotype largely confined to stem and progenitor compartments, without overt lineage collapse. This specificity may reflect cooperation between YY1 and other hematopoietic transcription factors or chromatin regulators that target PcG complexes to distinct genomic loci in HSCs. Identifying these YY1 PcG target genes and defining how their repression is dynamically regulated during aging will be important directions for future studies.

In summary, we establish YY1 PcG activity as a fundamental epigenetic mechanism that preserves HSC quiescence, metabolic restraint, and long-term self-renewal. Loss of this function drives premature HSC aging characterized by myeloid bias, metabolic activation, and functional exhaustion. These findings not only advance our understanding of the epigenetic regulation of stem cell aging but also suggest that restoring or stabilizing PcG-dependent repression may represent a potential strategy to rejuvenate aged HSCs and improve hematopoietic function in aging and disease.

## Methods

### Mice

*Yy1^+/ΔREPO^* mice were generated using CRISPR/Cas9 method. CRISPR tools were used to design two sgRNAs (5’ - CACCGGACCCTGGAGGGCGAGTTCT- 3’ and 3’CCTGGGACCTCCCGCTCAAGACAAA - 5’) flanking the REPO region. These sgRNAs were microinjected into fertilized C57BL/6J zygotes together with in vitro-transcribed Cas9 mRNA. Zygotes were cultured to the 2-cell stage and implanted into pseudo pregnant females at the Transgenic and Chimeric Mouse Facility at the University of Pennsylvania. Founder (F0) mice were genotyped by PCR and confirmed using Sanger sequencing. F0 mice were bred with wild-type C57BL/6 to achieve germline transmission, and F1 progeny were genotyped. Mice were subsequently backcrossed with C57BL/6 for 9 generations. For conditional deletion of the YY1 REPO domain, *Yy1^+/ΔREPO^* mice were crossed with *Yy1^f/f^* mice to generate *Yy1^f/ΔREPO^* mice. *Yy1^f/ΔREPO^* were crossed with *Yy1^f/+^ Mx1-Cre* or *Yy1^f/+^ Vav-Cre* mice to generate *Yy1^f/ΔREPO^ Mx1-Cre* or *Yy1^f/ΔREPO^ Vav-Cre* mice respectively. *Yy1^f/+^ Mx1-Cre / Vav-cre* heterozygous mice were generated by crossing *Mx1-Cre / Vav-cre* mice with *Yy1^f/f^* mice. To induce Cre recombinase expression in mice with *Mx1-Cre* background, 7- to 12-week-old mice were injected with 100 μg of polyinosinic-polycytidylic acid (pI-pC) every other day for 4 doses and all experiments were performed 7 days post the last pI-pC injection. The *Vav* promoter is activated at embryonic day 11.5 and drives Cre recombinase expression specifically in the hematopoietic system (53). C57BL/6 mice were used for experiments involving young (7 – 12 weeks old) and aged (24 months old) mice. All experimental protocols involving mice were approved by the Institutional Laboratory Animal Care and Use Committee of the University of Wisconsin-Madison and the University of Pennsylvania.

### Genotyping

Genotyping was performed by extracting DNA from tail samples of each mouse. PCR reactions were performed with the Platinum™ II Taq Hot-Start DNA Polymerase using respective primers to detect *Cre* allele, loxP-flanked *Yy1^f^* allele, and *Yy1^ΔREPO^*allele. The reaction cycles are as follows: Initial denaturation at 94°C for 2 minutes followed by: 34 cycles of denaturation at 94°C for 15 seconds, annealing at 60°C for 15 seconds, extension at 68°C for 15 seconds, and final extension at 68°C for 5 minutes.

### Western Blot

Whole cell lysates were prepared by lysing bone marrow cells in 4x SDS sample buffer and were loaded onto 10% Mini-PROTEAN® TGX™ Gels from Bio-Rad. Gels were run in 1x running buffer containing 25 mM Tris, 192 mM glycine, and 3.5 mM SDS. Proteins were transferred onto the nitrocellulose membrane using a transfer buffer containing 25 mM Tris, 201 mM glycine, 20% methanol by volume. Antibodies detecting target proteins YY1 (EPR4652, Abcam #ab109237) and Vinculin (E1E9V, Cell Signaling #13901) were used according to the manufacturer’s recommendation. Blots were developed using the enhanced chemiluminescence (ECL) system from Bio-Rad.

### Antibodies

Fluorophore or biotin-conjugated antibodies specific for the following surface antigens were purchased from eBioscience: CD3 (145-2C11), CD4 (RM4-5), CD8 (53-6.7), B220 (RA3-6B2), TER119 (TER-119), Gr-1 (RB6-8C5), IgM (eB121-15F9), CD19 (eBio1D3), IL-7Rα (A7R34), Mac1 (M1/70) and Thy1.2 (53-2.1), CD45.2 (104), CD45.1 (A20), Sca1 (D7), c-Kit (2B8), CD48 (HM48-1), CD34 (RAM34), CD61 (2C9.G3). CD135 (A2F10), CD16/32 (93) and CD150 (TC15-12F12.2) were purchased from BioLegend.

### Primary mouse evaluation

Bone marrow cells from *Yy1^+/+^*, *Yy1^+/−^*, and *Yy1^−/ΔREPO^* mice were harvested in phosphate-buffered saline (PBS) with 2 % fetal bovine serum (FBS) and passed through 70-μm cell strainers to obtain single-cell suspension. The cells were spun down and resuspended in PBS containing 2% FBS before antibody staining. . Lineage (Lin^−^) markers for HSPC evaluation include cocktail of biotin conjugated antibodies against CD3e, CD4, CD8, Ter119, B220, CD19, Gr1, IgM and IL-7Rα. Cells were gated as LSK (Lin^−^Sca1^+^c-kit^+^), LT-HSC (Lin^−^Sca1^+^c-kit^+^CD150^+^CD48^−^), ST-HSC (Lin^−^Sca1^+^c-kit^+^CD150^−^CD48^−^), MPP (Lin^−^Sca1^+^c-kit^+^CD150^−^CD48^+^), MP (Lin^−^Sca1^−^c-kit^+^), CMP (Lin^−^Sca1^+^c-kit^−^CD34^+^CD16/32^−^), GMP (Lin^−^Sca1^+^c-kit^−^CD34^+^CD16/32^+^) and MEP (Lin^−^Sca1^+^c-kit^−^CD34^−^CD16/32^−^) populations. DAPI (Thermo Fisher Scientific) was used to exclude non-viable cells. Flow cytometers LSR Fortessa II (BD Biosciences) or Aurora (Cytek) were used for data acquisition and analysis was performed using BD FlowJo v10.0.7 software.

### HSC Transplantation

For primary HSC transplantation, LT-HSCs (Lin^−^c-Kit^+^Sca1^+^CD150^+^CD48^−^) from 6- to 8-weekold *Mx1-Cre*, *Yy1^f/+^ Mx1-Cre*, and *Yy1^f/ΔREPO^ Mx1-Cre* CD45.2^+^ mice were sorted using BD FACSAria cell sorter (UW-Madison Carbon Cancer Center Flow Cytometry and Cell Sorting Facility) pre-pIpC injection. Per each mouse, 500 LT-HSCs were mixed with 0.5 million competitor cells from C57BL/6 (CD45.1^+^) mice and were injected retro-orbitally into lethally irradiated (8.5 Gy) C57BL/6 (CD45.1^+^) recipient mice. Mice were injected with pI-pC at 4 weeks post HSC transplantation to delete endogenous YY1. Peripheral blood chimerism was monitored every 4 weeks till 20 weeks post primary HSC transplant and bone marrow analysis and secondary transplant were performed at week 20. For secondary transplantation, 10 million bone marrow cells were harvested from primary transplant mice and were injected retro-orbitally into lethally irradiated C57BL/6 (CD45.1^+^) recipient mice. The chimeric recipient mice were monitored via peripheral blood analysis every 4 weeks for 16 weeks and bone marrow analysis was performed at week 16.

### Colony formation assay

Bone marrow cells from *Yy1^+/+^*, *Yy1^+/−^*, and *Yy1^−/ΔREPO^* mice were harvested and cultured in MethoCult™ GF M3434 media (StemCell™ Technologies) in 35mm dishes. BFU-E, CFU-GM and CFU-GEMM colonies were counted 12 – 14 days after plating.

### Ki67 proliferation assay

Bone marrow cells from C57BL/6 young, C57BL/6 aged, *Yy1^+/+^*, *Yy1^+/−^*, and *Yy1^−/ΔREPO^* mice were harvested and were stained for lineage markers and fixed with 4% paraformaldehyde. Cells were permeabilized using 0.1% saponin in PBS and were stained with PE-conjugated Ki67 (BD Biosciences, 1:40) and DAPI (Thermo Fisher Scientific, 1:1000). Data was acquired from LSR Fortessa (BD Biosciences) and analyzed using BD FlowJo v10.0.7 software.

### Seahorse assay

Mitochondrial respiration was assessed using the Seahorse XF Mito Stress Test (Agilent #103015-100) on an Agilent Seahorse XF Analyzer. Lin⁻c-Kit⁺ cells from C57BL/6 young, C57BL/6 aged, *Yy1^+/+^*, *Yy1^+/−^*, and *Yy1^−/ΔREPO^*mice were sorted and plated at 2 ×10⁵ cells/well on Cell-Tak-coated Seahorse XF microplates. Cells were incubated for 1 h at 37°C in non-CO₂ conditions in Seahorse XF Assay Medium supplemented with 1 mM pyruvate, 2 mM glutamine, 3 mg/mL glucose, 1× Pen-Strep, 100 ng/mL SCF, 10 ng/mL IL-6, and 5 ng/mL IL- 3. OCR was measured at baseline and after sequential injections of oligomycin (2 µM), FCCP (2 µM), and rotenone/antimycin A (0.5 µM each). Mitochondrial parameters, including basal respiration, ATP production, maximal respiration and spare respiratory capacity were calculated and normalized to cell number.

### Intracellular ROS Measurement

Bone marrow cells from C57BL/6 young, C57BL/6 aged, *Yy1^+/+^*, *Yy1^+/−^*, and *Yy1^−/ΔREPO^* mice were harvested and were stained for lineage markers. Cells were stained with 5 uM cloromethyl-2′,7′-dichlorofluorescin diacetate (CM-H2DCFDA, Thermo Fisher Scientific, #C6827) and were resuspended in DAPI (1:50,000). Upon oxidation, CM-H2DCFDA is converted to 2′, 7′-dichlorofluorescein (DCF), which can be detected by fluorescence spectroscopy (excitation/emission spectra of 495nm/529nm, respectively). ROS levels were quantified by using Cytek Aurora and BD FlowJo v10.0.7 software. Bone marrow cells without CM-H2DCFA treatment or treated with H_2_O_2_ and CM-H2DCFA were used as negative or positive control respectively.

### Senescence assay

CellEvent™ Senescence Green Flow Cytometry Assay Kit (ThermoFisher Scientific #C10841) which detects cellular senescence via β-galactosidase hydrolysis was used for this assay. Bone marrow cells from C57BL/6 young, C57BL/6 aged, *Yy1^+/+^*, *Yy1^+/−^*, and *Yy1^−/ΔREPO^* mice were harvested and were stained for lineage markers and fixed with 4% paraformaldehyde. Cells were then incubated with CellEvent™ Senescence Green Probe which contains two galactoside moieties specific to β-galactosidase. The enzyme cleaved product can be detected by fluorescence spectroscopy (excitation/emission spectra of 490nm/514nm, respectively). Fluorescent levels were quantified by using LSR Fortessa and BD FlowJo v10.0.7 software. Bone marrow cells without green probe treatment were used as negative control.

### RNA-seq data analysis

A total of 11 samples were chosen for RNA sequencing (RNA-Seq), 4 from *Yy1^+/+^*, 3 from *Yy1^+/−^*, and 4 from *Yy1^−/ΔREPO^*mice. Total RNA was purified from HSCs (Lin^−^c-Kit^+^Sca1^+^CD150^+/−^CD48^−^) sorted from all three cohorts using RNeasy Micro kit (Qiagen). Sequencing libraries were prepared by using Takara SMART-Seq V4 Ultra-Low Input according to the manufacturer’s specifications and were sequenced by an Illumina NovaSeq 6000 at the University of Wisconsin Biotechnology Center Gene Expression Center. RNA-seq reads were aligned by STAR (version 2.5.2b) to the mouse genome (version mm10) with GENCODE basic gene annotations (version M22). Gene expression levels were quantified by RSEM (version 1.3.0), and differential expression was analyzed by edgeR (version 3.36.0). A differentially expressed gene was required to have at least two-fold changes, an adjusted p-value < 0.05, and transcript per million (TPM) ≥ 1 in all the replicates in at least one of the two conditions in comparison. GSEA was performed by fgsea (version 1.20.0) with gene sets from the Molecular Signatures Database (version 7.1 downloaded from https://bioinf.wehi.edu.au/MSigDB/).

### Statistical analysis

All statistical analyses were performed using GraphPad Prism v7.04 software. Differences between the cohorts were determined using a student’s t-test when comparing two cohorts, or a one-way analysis of variance (ANOVA) followed by Tukey’s post-hoc test when comparing 3 or more cohorts. Two-way ANOVA followed by Tukey’s post hoc test was used to compare cells in different phases of the cell cycle. Two-way ANOVA was performed to determine the variation between line graphs. P-values ≤ 0.05 were considered statistically significant.

## Acknowledgments

We would like to acknowledge the University of Wisconsin Carbone Comprehensive Cancer Center (UWCCC) for providing access to their Shared Services, including the Flow Cytometry Core and Cancer Informatics Shared Resources, supported by UWCCC grant P30 CA014520. Additionally, this research was funded by grants TL1TR002375 and UL1TR002373, awarded to the University of Wisconsin-Madison Institute for Clinical and Translational Research (ICTR) through the National Center for Advancing Translational Sciences (NCATS). Further support for this work came from NIH grants R03OD026603 and R01HL146540 awarded to X. P., and NIH grant R01 AI155540 awarded to M. A.

## Author Contributions

X.P. designed experiments. S.S., J. L., Y.W., Y.L., C.H., and A.K. performed experiments. S. S., Y.W., P. L., Y.L., C.H., A.K., M.A., and X.P. analyzed and interpreted the data. S.S., P.L., M.A., L. R., and X.P. wrote the manuscript.

## Data Availability Statement

The datasets generated and/or analyzed during the current study are available from the corresponding author upon reasonable request.

