## Supplemental information for "YY1-Mediated Polycomb Group Function Safeguards Hematopoietic Stem Cells from Premature Aging"

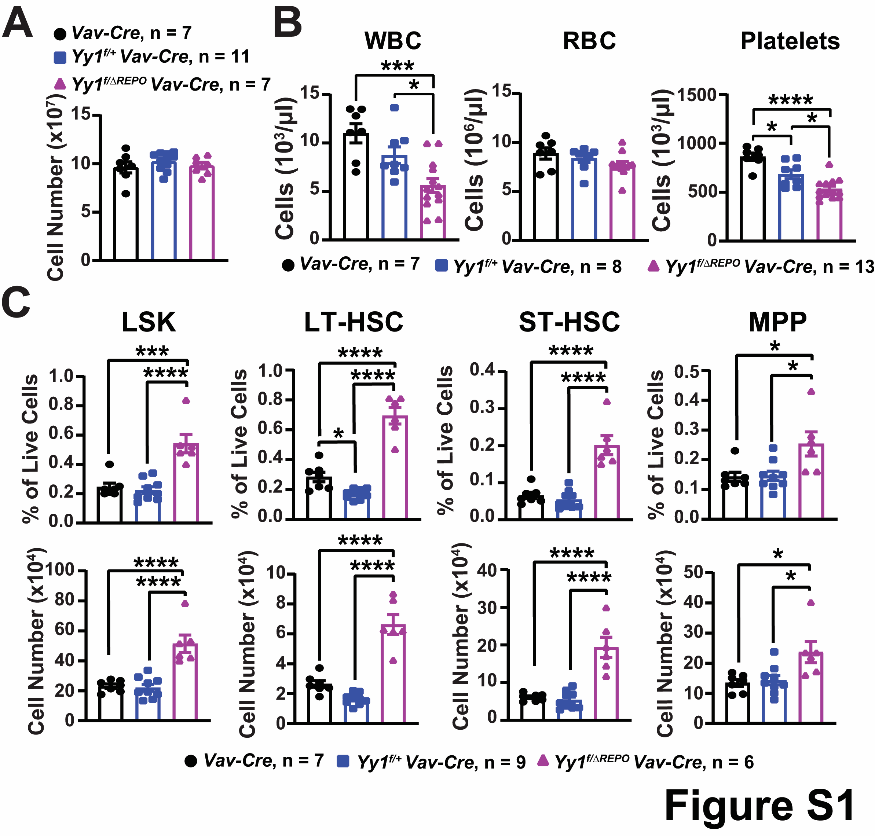

**Supplemental Figure 1**: **Conditional deletion of the YY1 REPO domain leads to expansion of immunophenotypic HSCs.** (**A**) Total BM numbers. (**B**) Complete blood count analysis. (**C**) Quantification of percentage and absolute number of LSK, LT-HSC, ST-HSC, and MPP populations. N represents the number of mice; Data are presented as means ± SEM; *P < 0.05, ***P < 0.001, ****P < 0.0001.
